# Age-related changes in the neural networks supporting semantic cognition: A meta-analysis of 47 functional neuroimaging studies

**DOI:** 10.1101/167064

**Authors:** Paul Hoffman, Alexa M. Morcom

## Abstract

Semantic cognition is central to understanding of language and the world and, unlike many cognitive domains, is thought to show little age-related decline. We investigated age-related differences in the neural basis of this critical cognitive domain by performing an activation likelihood estimation (ALE) meta-analysis of functional neuroimaging studies comparing young and older people. On average, young people outperformed their older counterparts during semantic tasks. Overall, both age groups activated similar left-lateralised regions. However, older adults displayed less activation than young people in some elements of the typical left-hemisphere semantic network, including inferior prefrontal, posterior temporal and inferior parietal cortex. They also showed greater activation in right frontal and parietal regions, particularly those held to be involved in domain-general controlled processing, and principally when they performed more poorly than the young. Thus, semantic processing in later life is associated with a shift from semantic-specific to domain-general neural resources, consistent with the theory of neural dedifferentiation, and a performance-related reduction in prefrontal lateralisation, which may reflect a response to increased task demands.

## Introduction

Semantic knowledge, of the meanings of words and properties of objects, shapes our understanding of the world and guides our behaviour. Most of our interactions with the environment, linguistic and non-linguistic, require us to harness this knowledge in some way. This use of semantic knowledge is often termed semantic cognition (Rogers and McClelland, 2004). Unsurprisingly, given its central role in higher cognitive function, semantic cognition activates a complex set of brain regions which overlap with other neural systems such as the multiple demand network (Duncan, 2010) and the default mode network (Buckner et al., 2008). In this meta-analysis, we investigated age-related differences in the functional neuroanatomy of semantic cognition. While formal meta-analysis techniques have been used to investigate functional brain activation in a number of domains (Li et al., 2015; Maillet and Rajah, 2014; Spreng et al., 2010), this is the first to focus on semantic cognition specifically. This is important because most aspects of semantic processing are thought to remain stable into older age, in stark contrast to the declines in function observed in many other cognitive domains (Nilsson, 2003; Nyberg et al., 1996; Park et al., 2002; Rönnlund et al., 2005; Salthouse, 2004; Verhaeghen, 2003). Important insights into the nature of successful cognitive ageing can be gained through better understanding of the changes in neural activity that underlie this maintenance of function. In what follows, we first provide an overview of the neural correlates of semantic cognition, as revealed by studies of young people. We then consider the predictions made by current theories of neurocognitive ageing for age-related differences in the networks engaged by semantic cognition in younger and older adults, before testing these predictions in a formal meta-analysis of 47 neuroimaging studies.

### The neural basis of semantic cognition

Semantic cognition activates a left-lateralised network in young adults, including frontal, temporal and parietal regions (Binder et al., 2009; Noonan et al., 2013). Key regions are illustrated in blue in Figure 1 (alongside other networks to be described later). The ventral anterior temporal lobe (vATL) is thought to be involved in the storage of multi-modal semantic representations (Lambon Ralph et al., 2017). This is based on the strong association between damage to this region and the clinical syndrome of semantic dementia, which involves a profound and selective loss of semantic knowledge (Patterson et al., 2007). fMRI studies often overlook vATL, in part because of well-known technical difficulties in acquiring signal from the ventral temporal cortices, due to the proximity of air-filled sinuses (Devlin et al., 2000). However, recent studies using methods that combat these issues have reliably identified activity in the left vATL during semantic processing (Halai et al., 2015; Hoffman et al., 2015; Humphreys et al., 2015).

**Figure 1:**
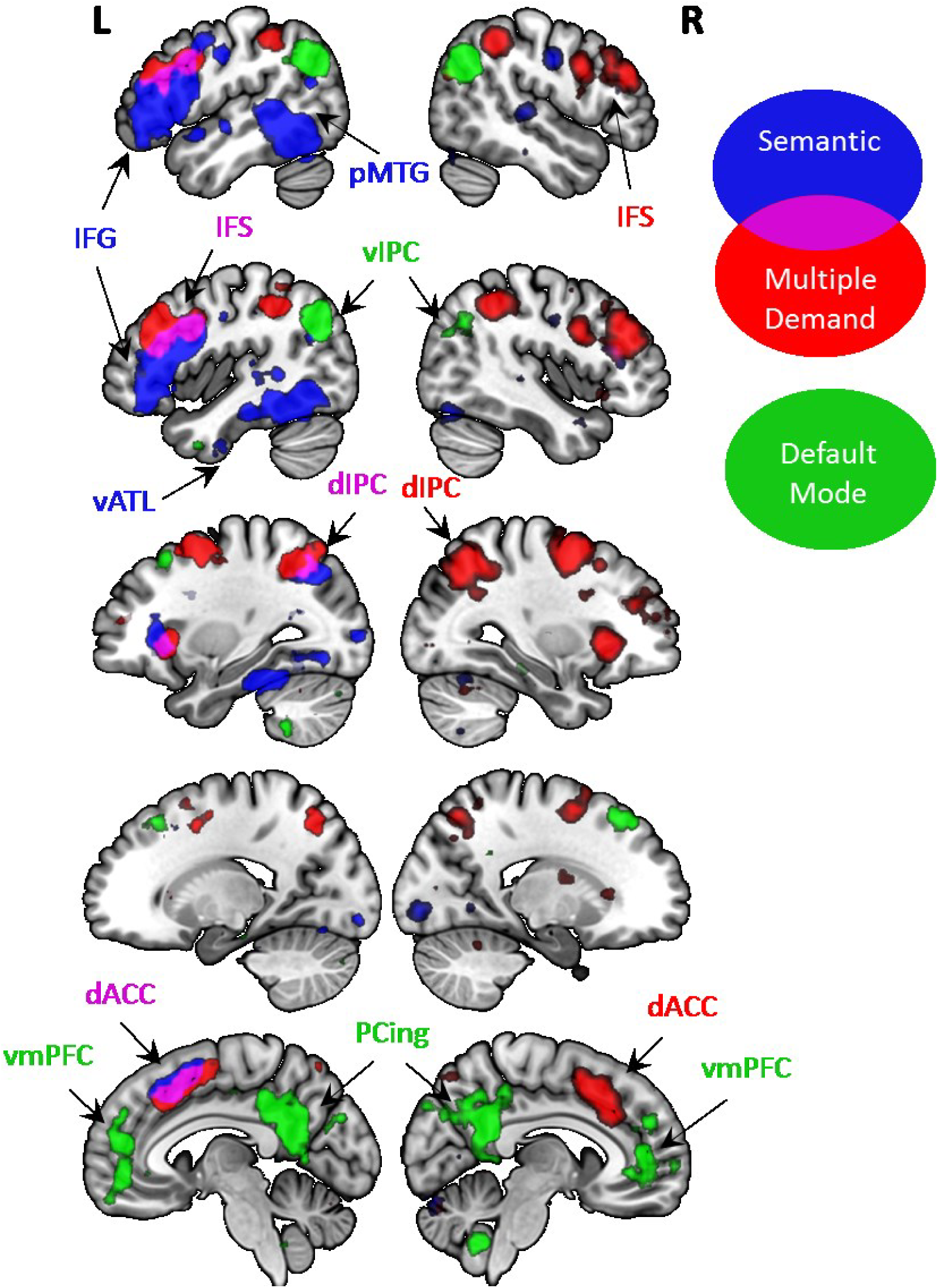
Regions typically associated with semantic processing and with the multiple demand and default mode networks. Figure show areas of activation associated with particular topics in the Neurosynth database of over 10,000 neuroimaging studies (Yarkoni et al., 2011). Topics were extracted using automated analysis of terms used in the target articles (Poldrack et al., 2012). The semantic topic included the keywords [semantic, words, meaning, picture, conceptual, association, knowledge]. The multiple demand topic included [task, performance, control, executive, difficulty, demands, goal]. The default mode topic included [network, resting, default, intrinsic, spontaneous]. The database does not discriminate between young and older participants; however, since the vast majority of neuroimaging participants are young, these networks predominately reflect activation patterns in young adults. IFG = inferior frontal gyrus; pMTG = posterior middle temporal gyrus; IFS = inferior frontal sulcus; vIPC = ventral inferior parietal cortex; vATL = ventral anterior temporal lobe; dIPC = dorsal inferior parietal cortex; dACC = dorsal anterior cingulate cortex; PCing = posterior cingulate cortex; vmPFC = ventromedial prefrontal cortex.

Other regions are involved in the executive regulation of semantic knowledge, ensuring that task and context-appropriate information is activated (Jefferies, 2013). This is critical because we store a wide range of knowledge about any concept and different aspects of this information are important in different situations. For example, the relevant semantic features of *pianos* change depending on whether one is asked to play a piano or to move one across the room (Saffran, 2000). This element of semantic processing, often termed semantic control, has chiefly been associated with activity in the left inferior frontal gyrus (IFG) (Badre and Wagner, 2002; Hoffman et al., 2010; Thompson-Schill et al., 1997). More recently, it has become clear that left posterior middle temporal gyrus (pMTG) is also activated by manipulations of semantic control (Noonan et al., 2013; Whitney et al., 2011). The two regions also display strong structural and functional interconnectivity (Turken and Dronkers, 2011). Current theories hold that both IFG and pMTG serve to regulate performance in semantic tasks by exerting top-down control over the activation of semantic representations in the vATL (Lambon Ralph et al., 2017).

Semantic tasks also activate some areas within the “multiple demand” network (MDN) (Duncan, 2010; Fedorenko et al., 2013). This network, sometimes termed the frontoparietal control system (Vincent et al., 2008), comprises a set of brain regions that respond to increasing task demands across many cognitive domains and are thought to be involved in the planning and regulation of goal-directed cognition and behaviour (see red regions in Figure 1). MDN regions activated during semantic processing include left dorsal inferior parietal cortex (dIPC; in the region of the intraparietal sulcus), left inferior frontal sulcus (IFS; superior to IFG) and the dorsal anterior cingulate (dACC; often including the pre-supplementary motor area) (Noonan et al., 2013). Importantly, however, MDN activity during semantic tasks is usually restricted to left-hemisphere structures, in contrast to other domains such as visuospatial processing, which preferentially activate right-hemisphere elements of this network (Shulman et al., 2010). Thus, although semantic tasks recruit elements of the domain-general MDN as well as semantic-specific brain regions, there is a bias in both cases towards left-hemisphere activation.

Finally, semantic processing has been linked with the default mode network (DMN), a set of brain regions that display greater activity during rest periods when participants are not engaged in an overt task (Buckner et al., 2008; Raichle et al., 2001). Core areas of the DMN include the ventral inferior parietal cortex (vIPC) bilaterally, the ventromedial prefrontal cortex (vmPFC) and the posterior cingulate (pCing) (see green regions in Figure 1). Some descriptions of the DMN include the ATL, which has also been strongly implicated in semantic representation (Buckner et al., 2008; Humphreys et al., 2015). Some researchers have proposed that DMN activation during rest is a consequence of implicit semantic processing, as participants at rest engage in daydreaming and other semantically-rich forms of self-directed thought in the absence of any external stimulus (Binder et al., 1999). However, other studies have shown that, with the exception of the ATL, DMN regions are not activated by explicit semantic tasks, suggesting that these regions are unlikely to make a major contribution to semantic cognition (Hoffman et al., 2015; Humphreys et al., 2015).

### Age-related changes in functional brain networks

In addition to well-known changes in brain structure (Raz et al., 2005), functional imaging studies have suggested that ageing may affect how brain networks are configured and how they respond to cognitive challenges. Although relatively few neuroimaging studies of cognitive ageing have been concerned with semantic cognition specifically, two main general principles of functional reorganisation have been proposed (Grady, 2012; Morcom and Johnson, 2015). Each outlines specific regional patterns of age-related differences which are frequently interpreted in terms of compensatory shifts which help to support performance. According to alternative views, increases in activation or activation of additional regions in ageing may reflect reduced specificity of neuronal responses rather than compensation. This may be due to noisy neuronal representations (Li et al., 2001) or impaired ability to regulate activity across networks (Grady et al., 1994; Logan et al., 2002). It is difficult to adjudicate between these mechanisms using functional imaging measures of activation (Lövdén et al., 2010; Morcom & Johnson, 2015). In this meta-analysis, we were interested in the proposed principles of reorganisation and the predictions they make for age-related differences in the networks supporting semantic function.

One long-standing observation is that older adults often show more activation in visual processing tasks than young adults in prefrontal cortices and may also show less activation in occipitotemporal cortices (Davis et al., 2008; Grady et al., 1994; Maillet and Rajah, 2014; Spreng et al., 2010)although increased activation has also been observed in posterior cortical regions in older adults, (e.g., Grady et al., 1994). This pattern, termed PASA (posterior-to-anterior shift in aging; Dennis and Cabeza, 2008) is proposed to reflect an upregulation in the executive control processes supported by the prefrontal cortices, to compensate for less efficient visual processing. Since most studies of semantic processing involve presentation of visual stimuli (either words or pictures), a straight-forward prediction of the PASA theory is that older adults will exhibit increased prefrontal activation, and reduced visual cortex activation, during semantic tasks. As the left IFG is strongly implicated in executive regulation of semantic knowledge, this is a possible site for such an upregulation. Alternatively, or in addition, the increased demands may cause older adults to recruit MDN regions, which respond to increased demands in semantic processing as well as in other cognitive domains.

In parallel, researchers have frequently noted age-related reductions in the laterality of prefrontal activation, with tasks that elicit lateralised activity in young people displaying a more bilateral pattern in older adults. This phenomenon, termed HAROLD (hemispheric asymmetry reduction in older adults; Cabeza, 2002), has, like PASA, been proposed to reflect a compensatory response (Grady, 2012). Indeed, in a meta-analysis of 80 neuroimaging studies using a range of cognitive tasks, increased recruitment of right prefrontal regions in older adults was only observed where older people performed more poorly than their young counterparts (Spreng et al., 2010). This is compatible with the view that the increased recruitment helps to maintain performance under difficult conditions (but also with the possibility that increasing task demand triggers or enhances nonspecific responses; Logan et al., 2002). When performance was equivalent there was no evidence of HAROLD: instead, older adults engaged left dorsolateral PFC more and left IFG less than the young, consistent with greater use of MDN resources (Spreng et al., 2010).

In semantic tasks, IFG activation is strongly left-lateralised in young adults, with the right IFG only called upon to contribute under the most demanding conditions (Krieger-Redwood et al., 2015; Noonan et al., 2013). If semantic tasks become more difficult in older people, one might expect them to engage this region more frequently, resulting in a HAROLD pattern. This hypothesis is consistent with a further proposal that the recruitment of brain regions is governed by a load-dependent function that shifts in older age, the CRUNCH theory (compensation-related utilization of neural circuits hypothesis; Reuter-Lorenz and Cappell, 2008). CRUNCH states that older people tend to increase their recruitment of neural resources at a lower level of task demand than young people, in order to maintain performance at a similar level (see also Park and Reuter-Lorenz, 2009). It also states that additional recruitment of brain regions is subject to a ceiling effect as task demand increases, and after this point young people display greater activation. Left IFG is one region where this may be a likely outcome. This region displays robust activation in young adults for almost all semantic tasks and thus may have little spare capacity for additional recruitment in later life. Of course, these predictions assume that older people find semantic tasks more demanding than young people. While this assumption is uncontroversial for many areas of cognition, it is less certain in the semantic tasks, on which young and old often perform at similar levels (Nilsson, 2003; Nyberg et al., 1996; Park et al., 2002; Rönnlund et al., 2005; Salthouse, 2004; Verhaeghen, 2003).

Finally, older adults also frequently display increased activation of the DMN (Grady et al., 2010; Persson et al., 2007). In most cases, however, this is unlikely to reflect an adaptive compensatory strategy, since activity decreases rather than increases in DMN regions are associated with successful completion of most tasks (Buckner et al., 2008; Persson et al., 2007). Age-related differences in this network may therefore indicate a failure in older adults to deactivate neural systems that are unrelated to the task at hand. The failure to inhibit DMN activity during demanding tasks may be an example of the broader phenomenon of dedifferentiation of neural activity in later life (Grady, 2012).

To investigate age-related differences in the neural basis of semantic cognition, we performed an activation likelihood estimation (ALE) meta-analysis of 47 functional neuroimaging studies that contrasted young and older adults on tasks involving semantic processing. Theories of neurocognitive ageing posit that age-related changes in activation are either a cause of or response to diminished task performance in older people. To assess whether performance declined with age in the studies we analysed, we computed behavioural effect sizes for the difference between young and older participants wherever possible. This allowed us to divide studies into those in which young and older participants were well-matched in performance and those in which young people outperformed older people, allowing us to investigate whether these two situations led to different outcomes.

## Analysis Method

### Study selection

We searched for peer-reviewed studies published between January 1990 and August 2016, in which young and older adults were compared on tasks that required semantic processing. An initial search was conducted on 25^th^ August 2016 using the Scopus database for articles containing the following terms in their title or abstract: (fMRI OR PET OR neuroimaging) AND (age OR ageing OR ageing OR older) AND (semantic* OR speech OR language OR comprehension OR fluency OR naming OR sentence*). This yielded 1176 studies, which were screened for inclusion in the meta-analysis. Further candidate studies were identified by searching the reference lists of studies that passed the screening process, and those of previous meta-analyses of functional neuroimaging studies of cognitive ageing (Li et al., 2015; Maillet and Rajah, 2014; Spreng et al., 2010).

Inclusion criteria were as follows:

1. Experimental paradigm contrasted two conditions, one of which had a greater involvement of semantic processing. A broad definition of semantic knowledge was used, which included the meanings of words and sentences as well as knowledge relating to meaningful objects or faces. Tasks included explicit semantic decisions (e.g., animacy or concreteness judgements), tasks that implicitly engage semantic processing (e.g., lexical decision or passive listening to speech) and semantically-driven word retrieval tasks (e.g., category fluency and naming). Stimuli were most often written words, although some studies presented spoken words, pictures or familiar odours. There were also a number of studies whose main focus was episodic memory but which used semantic judgements as an incidental memory encoding task (e.g., Madden et al., 1999). These studies were included if they reported activations elicited by the semantic encoding phase independent of later retrieval activity. We excluded any studies using only the subsequent memory paradigm (e.g., Morcom et al., 2003).
2. Study included a healthy young adult (mean age < 45) and older adult (mean age > 60) group and reported whole-brain activation peaks either from each group independently or from contrasts of the two groups. In addition, a small number of studies were included that reported positive and negative effects of ageing using a parametric design, with participants spanning from young to older age.

A total of 47 studies met the inclusion criteria (see Table 1). A number of otherwise eligible studies could not be included, either because they presented activation maps visually but did not report peak activation co-ordinates (e.g., Logan et al., 2002), because they only reported deactivations relative to rest (e.g., Persson et al., 2007) or because analyses discriminating between young and older adults were only performed in regions of interest (e.g., Shafto et al., 2010).

**Table 1:** Details of studies included in the meta-analysis

| Study # | First author | Year | Semantic task | Baseline task | Effect size (Cohen's d) | Imaging modality | Number of peaks |  |  |  | Young group |  | Older group |  |
| --- | --- | --- | --- | --- | --- | --- | --- | --- | --- | --- | --- | --- | --- | --- |
|  |  |  |  |  |  |  | O | Y | O>Y | Y>O | N | mean age | N | mean age |
| 1 | Anderson | 2000 | Visualise relationship between two words | Cued episodic recall of words | NA | PET |  |  | 11 | 13 | 12 | 24.4 | 12 | 68.5 |
| 2 | Baciu | 2016 | Category fluency & Picture naming & Picture semantic association | Rest & Shape naming & Shape matching | 1.67 & -0.32 & 0.50 | fMRI |  |  | 41 | 2 | 16 | 42.6 | 14 | 72.2 |
| 3 | Backman | 1997 | Spoken word stem completion | Passive viewing of word stems | NA | PET | 2 | 2 |  |  | 7 | 24.3 | 7 | 63.4 |
| 4 | Bergerbest | 2009 | Concreteness decisions on novel words | Concreteness decisions on old (primed) words | NA | fMRI | 7 | 3 | 11 | 2 | 16 | 23.4 | 15 | 78.7 |
| 5 | Berlingeri | 2010 | Picture naming+animacy decision & sentence plausibility judgements | Scrambled picture decision & old/new recognition judgements | 0.20 | fMRI |  |  | 10 | 19 | 24 | 26.5 | 24 | 62.0 |
| 6 | Cabeza | 1997 | Find meaningful relationship between two words | Episodic memory for word pairs | NA | PET |  |  | 10 | 11 | 12 | 26.0 | 12 | 70.0 |
| 7 | Daselaar | 2003a | Pleasantness decisions on written words | Motor response to arrow cues | -0.35 | fMRI | 3 | 4 |  | 1 | 20 | 32.7 | 21 | 66.2 |
| 8 | Daselaar | 2003b | Animacy decision on written words | Case judgement to written words | 0.38 | fMRI | 17 | 18 |  | 6 | 26 | 32.4 | 39 | 66.3 |
| 9 | Daselaar | 2005 | Word stem completion for novel words | Word stem completion to old (primed) words | -0.76 | fMRI | 3 | 5 |  | 3 | 25 | 32.5 | 38 | 66.4 |
| 10 | Davis | 2014 | Plausibility judgements on auditory sentences | Listening to auditory noise | 0.22 | fMRI |  |  | 2 |  | 50 | Parametric |  |  |
| 11 | Destrieux | 2012 | Category fluency | Rest | 1.02 | fMRI |  |  | 7 | 1 | 22 | 25.2 | 21 | 80.2 |
| 12 | Diaz | 2014 | Property verification on pictures | Perceptual and phonological judgements on pictures | 0.65 | fMRI |  |  | 30 | 4 | 16 | 23.5 | 16 | 68.2 |
| 13 | Donix | 2010 | Familiarity judgements on familiar faces | Familiarity judgements on unfamiliar faces | NA | fMRI | 9 | 7 | 1 |  | 12 | 30.4 | 12 | 62.1 |
| 14 | Eckert | 2008 | Repetition of words presented in noise | Words not recognised | 0.54 | fMRI |  |  | 4 |  | 15 | Parametric |  |  |
| 15 | Geva | 2012 | Rhyme judgements on pictures and words | Perceptual judgements on symbols and scrambled pictures | 0.09 | fMRI |  |  | 2 |  | 12 | 24.6 | 19 | 64.1 |
| 16 | Gold | 2009 | Lexical decisions on written words | Rest | <0† | fMRI |  |  | 3 | 4 | 15 | 22.9 | 14 | 74.7 |
| 17 | Grady | 1999 | Animacy decision on written words and pictures | Episodic encoding or perceptual decisions on written words and pictures | NA | PET |  |  |  | 8 | 12 | 23.0 | 12 | 66.2 |
| 18 | Grossman | 2002 | Comprehension judgements on written sentences | Perceptual task on pseudofont sentences | 0.56 | fMRI | 25 | 19 | 15 | 8 | 13 | 22.6 | 11 | 63.5 |
| 19 | Hwang | 2007 | Passive listening to spoken text | Rest | NA | fMRI | 7 | 7 |  |  | 12 | 26.0 | 12 | 70.0 |
| 20 | Iidaka | 2001 | Find meaningful relationship between two pictures | Form association between two abstract shapes | NA | fMRI | 10 | 11 |  | 2 | 7 | 25.7 | 7 | 66.2 |
| 21 | Johnson | 2001 | Relatedness judgements on spoken word pairs | Similarity decision on spoken nonwords | 0.18 | fMRI | 3 | 8 |  | 2 | 9 | 31.9 | 9 | 72.7 |
| 22 | Kalenzaga | 2015 | Imagine scene based on written word cue | Rest | NA | fMRI |  |  | 5 |  | 19 | 29.2 | 16 | 68.3 |
| 23 | Kareken | 2003 | Odour identification (matching smell to object) | Odour smelling | 0.82 | PET |  |  | 3 | 6 | 5 | 27.8 | 6 | 71.0 |
| 24 | Kounios | 2003 | Pleasantness decisions on written words | Judgements on nonwords | NA | fMRI | 8 | 3 | 2 | 2 | 16 | 23.4 | 16 | 73.9 |
| 25 | Kuchinsky | 2012 | Auditory word recognition | Listening to noise | 0.43 | fMRI |  |  | 6 |  | 36 | Parametric |  |  |
| 26 | Leshikar | 2010 | View two objects and generate a sentence containing both | View abstract patterns | NA | fMRI |  |  | 53 | 2 | 19 | 20.9 | 18 | 65.7 |
| 27 | Madden | 1996 | Lexical decisions on written words | Motor response on all words and nonwords | NA | PET | 3 | 5 |  | 4 | 10 | 22.5 | 10 | 68.2 |
| 28 | Madden | 1999 | Animacy decision on written words | Case judgement to written words | NA | PET | 5 |  | 2 |  | 12 | 23.2 | 12 | 71.0 |
| 29 | Madden | 2002 | Lexical decisions on written words | Perceptual task on consonant strings | NA | PET | 5 | 7 | 1 | 1 | 12 | 23.6 | 12 | 65.0 |
| 30 | Madden | 2010 | Size & animacy decisions on written words | Rest | 0.49 | fMRI |  |  | 17 | 2 | 20 | 22.4 | 20 | 69.6 |
| 31 | Maguire | 2003 | Truth judgements to auditory facts | Syllable judgments to jumbled word strings | 0.43 | fMRI | 9 | 18 |  |  | 12 | 32.4 | 12 | 74.8 |
| 32 | Marsolais | 2014 | Category fluency | Reciting months of the year | 0.43 | fMRI | 39 | 69 |  |  | 14 | 24.0 | 14 | 63.5 |
| 33 | Martins | 2014 | Word sorting by semantic category | Word sorting by rhyme or initial phoneme | NA | fMRI | 4 | 40 | 15 | 30 | 28 | 26.0 | 14 | 63.0 |
| 34 | McGeown | 2009 | Semantic association task on written words | Perceptual task on nonwords | 1.16 | fMRI | 4 | 11 | 3 | 17 | 10 | 23.1 | 9 | 75.1 |
| 35 | Meinzer | 2009 | Category fluency | Repeating the word "pause" | 3.14 | fMRI | 14 | 9 | 5 |  | 16 | 26.1 | 16 | 69.3 |
| 36 | Meinzer | 2012a | Category fluency | Repeating the word "rest" | 0.88 | fMRI | 12 | 14 | 10 |  | 14 | 24.6 | 14 | 69.2 |
| 37 | Meinzer | 2012b | Category fluency | Repeating the word "rest" | 1.14 | fMRI | 17 | 37 |  |  | 16 | 24.0 | 16 | 68.9 |
| 38 | Murty | 2008 | Indoor/outdoor decisions on photographs | Rest | 0.46 | fMRI |  |  | 12 | 9 | 30 | 25.6 | 30 | 61.2 |
| 39 | Neilson | 2006 | Familiarity judgements on names of famous people | Familiarity judgements on names of non-famous people | 0.42 | fMRI | 23 | 5 | 15 |  | 15 | 23.6 | 15 | 70.4 |
| 40 | Peelle | 2013 | Semantic feature matching on written words | Rest | -0.19 | fMRI |  |  | 15 |  | 18 | 24.4 | 21 | 65.0 |
| 41 | Roxbury | 2016 | Lexical decisions on spoken words | Lexical decisions on spoken nonwords | 0.58 | fMRI |  |  | 4 |  | 17 | 27.4 | 17 | 71.0 |
| 42 | Shafto | 2012 | Lexical decisions on spoken words | Pitch decisions on auditory stimuli | NA | fMRI | 6 | 20 |  |  | 14 | 23.9 | 16 | 75.8 |
| 43 | Stebbins | 2002 | Concreteness decisions on written words | Case judgement to written words | 0.10 | fMRI | 5 | 14 |  |  | 15 | 25.3 | 15 | 76.5 |
| 44 | Suzuki | 2001 | Odour identification task | Motor response with no odour | NA | fMRI | 2 | 15 |  |  | 6 | 30.0 | 6 | 73.0 |
| 45 | Tyler | 2010 | Monitoring auditory word strings for particular words | Monitoring auditory noise | NA | fMRI | 37 | 20 |  |  | 14 | 23.9 | 44 | 67.4 |
| 46 | Wierenga | 2008 | Picture naming | Viewing abstract shapes | 0.59 | fMRI | 7 | 3 | 16 |  | 20 | 25.1 | 20 | 74.9 |
| 47 | Wong | 2009 | Matching spoken words to pictures | Rest | 1.94 | fMRI |  |  | 7 | 4 | 12 | 21.8 | 12 | 67.5 |
|  |  |  |  |  |  | **TOTAL** | **286** | **374** | **338** | **163** | **723** | **26.0** | **766** | **69.1** |

Positive effect sizes indicate a performance advantage favouring young participants and negative sizes favouring older participants. NA = data not available. † indicates that there were insufficient data to compute an exact effect size but that the mean performance of older participants exceeded that of the young.

To classify studies based on performance differences, we calculated an effect size (Cohen’s d) for the difference between young and older participants, using a similar approach to Li et al. (2015). Effect sizes were computed based on the means and standard deviations of the two groups or from test statistics comparing the groups. Effect sizes were computed from number of correct responses/errors but not from reaction times, since older people exhibit general reductions in processing speed that may not reflect changes in semantic processing per se. The only exception to this rule was for two studies that required participants to make subjective judgements about concepts (pleasantness judgements; Daselaar et al., 2003; Grossman et al., 2002). Since it is not possible to score such judgements for accuracy, we used reaction time data for these studies. In both cases, responses were faster for the older group, so the effect could not be attributed to age-related slowing.

### ALE analyses

A series of Activation Likelihood Estimation (ALE) analyses were carried out using GingerALE 2.3.6 (Eickhoff et al., 2012; Eickhoff et al., 2009). This software takes activation peaks from neuroimaging contrasts of interest, across a range of independent studies, models the spatial distribution of these peaks and computes whole-brain activation likelihood maps. These maps can then be subjected to voxel-wise statistical tests to identify regions that are reliably activated across studies.

We used the ALE method to investigate regions activated by semantic processing in young and older adults and to explore age-related differences in these networks. We considered four types of activation foci, which we labelled Y, O, Y>O and O>Y (see Table 2 for numbers of peaks in each study type). Y and O refer to peaks obtained in independent analyses of each age group while Y>O and O>Y refers to peaks obtained in within-study contrasts of the two age groups. We analysed these four types separately because they give complementary information about the underlying neural networks. The Y and O peaks provide essential information about the spatial distribution of activation in each age group, allowing us to determine the degree to which the age groups activate similar networks during semantic processing. The within-study contrasts (Y>O and O>Y) provide information about differences in the degree to which each group activates specific regions.

**Table 2:** Number of studies and number of peaks available for each analysis

|  |  | Peak Type |  |  |  |
| --- | --- | --- | --- | --- | --- |
|  |  | Y | O | Y>O | O>Y |
| Number of studies contributing peaks | TOTAL | 26 | 27 | 25 | 31 |
|  | Performance-Equivalent |  |  | 8 | 9 |
|  | Performance-Reduced |  |  | 8 | 14 |
| Number of peaks | TOTAL | 374 | 286 | 163 | 338 |
|  | Performance-Equivalent |  |  | 23 | 71 |
|  | Performance-Reduced |  |  | 50 | 156 |

We conducted the following analyses:

1. Activation in young and older adults. These analyses considered Y and O peaks separately and identified areas consistently activated by each age group. A conjunction analysis was also performed to identify regions commonly activated by both groups.
2. Contrasts of young and older adults. These analyses used the Y>O and O>Y peaks to identify areas in which older adults reliably exhibited more or less activation than young adults. We also performed a laterality analysis for these ALE maps (following Rice et al., 2015b; Turkeltaub and Coslett, 2010). A mirror image of each ALE map was generated and subtraction analyses were performed to identify regions in which ALE values in one hemisphere were significantly higher than in the homologous region in the opposite hemisphere. This allowed us to formally test the lateralisation of activation differences.
3. Division of studies by behavioural effects. Finally, we formed two subsets of studies based on the effect sizes of the behavioural differences between the two age groups. We arranged all the studies in order of effect size and performed a median split. In the half with the smaller effect sizes, performance did not differ between young and older participants (Performance-Equivalent studies), while in the half with the larger effect sizes there was a performance difference favouring the young (Performance-Reduced studies). We performed separate ALE analyses of Y>O and O>Y peaks for these subsets of studies, to investigate the effect of behavioural performance on neural activity differences.

In GingerALE, each activation peak is modelled as a probability distribution centred on the peak co-ordinates, generated by Gaussian smoothing. This accounts for uncertainty in the true focus of activation due to between-subject variability. The full-width half maximum (FWHM) of the smoothing kernel is determined by the number of subjects generating the peak, and is based on estimates of between-subject variability in activity elicited in motor cortex by finger-tapping (Eickhoff et al., 2009). For the present analyses, we added 10mm to the smoothing kernel to account for the greater between-subject variability associated with higher cognitive functions (including semantic processing; see Tahmasebi et al., 2011). We used Turkeltaub et al.’s (2012) non-additive version of the ALE algorithm, which limits the influence of a single study reporting multiple peaks very close to one another. Peaks reported in Talairach space were converted to MNI space using the tal2icbm_spm transform (Lancaster et al., 2007). Analyses were thresholded using a permutation-based method for cluster-level inference (Eickhoff et al., 2012). A family-wise error cluster-corrected threshold of *p* < 0.05 was adopted (with a cluster-forming threshold of *p* < 0.01). For some analyses, the minimum cluster size indicated using this method was rather large (over 20,000mm^3^). To determine whether smaller clusters were present below the cluster-corrected threshold, we re-ran analyses with an uncorrected threshold of *p* < 0.01 and an arbitrary extent threshold of 1000mm^3^. Because these results were not corrected for multiple comparisons, we draw no strong inferences from them; however, they are provided as Supplementary Materials and we note where they are consistent with prior hypotheses about age-related effects.

The laterality analysis of Y>O and O>Y maps involved a subtraction of ALE maps. Cluster-level inference is not currently available for subtraction analyses so we instead initial individual analyses of each dataset were performed at *p* < 0.01, i.e., the same voxel threshold as in the main analyses and then adopted an uncorrected threshold of *p* < 0.05 (with a minimum cluster size of 500mm^3^) for the final subtraction map. All thresholds were computed using 5000 random permutations of the dataset.

## Results

Forty-seven studies were included in the meta-analysis, comprising a total of 723 young and 766 older participants (see Table 1). The mean age of young participants was 26.0 years (SD=4.1) and the mean age of older participants was 69.1 (SD=4.7). Table 2 shows the number of studies contributing Y, O, Y>O and O>Y peaks to the analyses reported below.

### Activation in young and older adults

Figure 2 shows ALE maps generated from separate analyses of young and older participants (using all Y and O peaks), as well as their overlap. Peak areas of convergence are reported for the separate analyses in Table 3 and for their conjunction in Table 4. Very similar regions were identified in the two populations. The overlap included several regions implicated in semantic processing, such as left IFG, left pMTG and dACC, as well as overlapping clusters in right IFG. The uncorrected analysis also revealed overlapping activation in dIPC (see Supplementary Figure 1), though this was not present at the cluster-corrected threshold.

**Figure 2:**
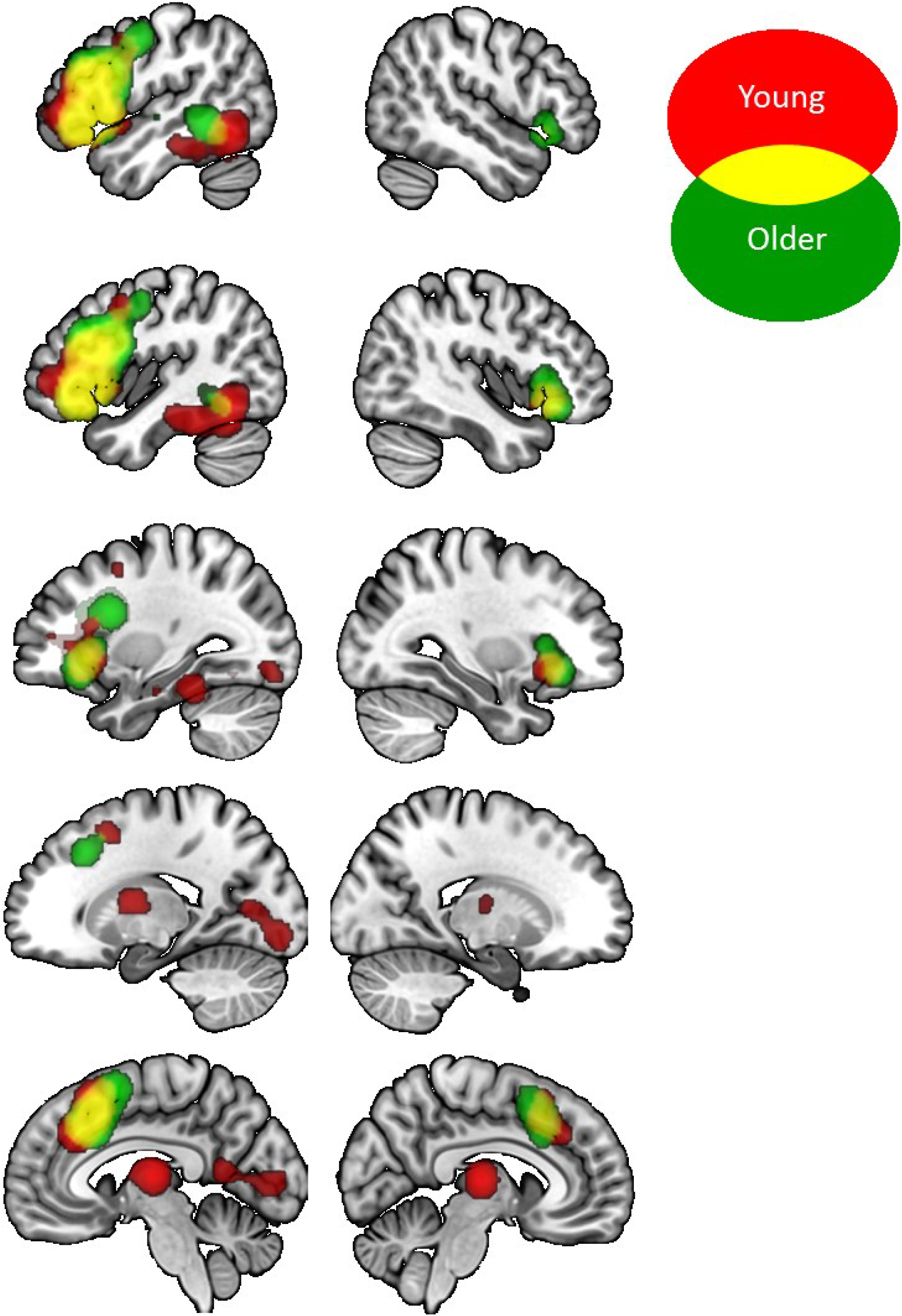
Activation likelihood maps for separate analyses of young and older people. Results are presented at a threshold of *p* < 0.05, corrected for multiple comparisons at the cluster level.

**Table 3:** ALE clusters for activation by young and older adults across all studies

| Cluster | Anatomical region | Volume (mm<sup>3</sup>) | BA | x | y | z | ALE value |
| --- | --- | --- | --- | --- | --- | --- | --- |
| *Young* |  |  |  |  |  |  |  |
| 1 | L lateral prefrontal | 55680 |  |  |  |  |  |
|  | L IFG (pars triangularis) |  | 47/45 | -50 | 26 | -2 | .0079 |
|  | L IFG (pars opercularis) |  | 44/45 | -50 | 22 | 18 | .0070 |
|  | L IFS/middle frontal gyrus |  | 6 | -44 | 4 | 44 | .0029 |
| 2 | L medial frontal & cingulate | 20832 |  |  |  |  |  |
|  | SFG/dACC |  | 8 | -4 | 20 | 48 | .0060 |
|  | SFG/ dACC |  | 32 | -4 | 22 | 44 | .0060 |
|  | SFG/ dACC |  | 24 | -4 | 28 | 36 | .0055 |
|  | L SFG |  | 8 | -24 | 6 | 50 | .0029 |
| 3 | L posterior temporal & occipitotemporal | 19512 |  |  |  |  |  |
|  | L inferior temporal gyrus |  | 37 | -46 | -60 | -10 | .0045 |
|  | L fusiform gyrus |  | 37 | -40 | -40 | -22 | .0036 |
|  | L parahippocampal /hippocampus |  | 20 | -32 | -16 | -22 | .0025 |
| 4 | Thalamus & L caudate | 13392 |  |  |  |  |  |
|  | Thalamus |  | - | -2 | -16 | 8 | .0050 |
|  | Thalamus |  | - | -12 | -6 | 12 | .0037 |
|  | L caudate |  | - | -12 | -2 | 12 | .0037 |
| 5 | L occipital lobe | 10328 |  |  |  |  |  |
|  | L lingual gyrus |  | 17 | -12 | -78 | 4 | .0032 |
|  | L lingual gyrus |  | 18 | -26 | -86 | -12 | .0031 |
|  | L occipital pole |  | 18 | -16 | -90 | -6 | .0029 |
|  | L precuneus |  | 17 | -10 | -58 | 8 | .0028 |
|  | L precuneus |  | 17 | -6 | -58 | 12 | .0028 |
| 6 | R IFG | 6664 |  |  |  |  |  |
|  | R anterior IFG (pars orbitalis) |  | 47 | 38 | 24 | -8 | .0047 |
| *Older* |  |  |  |  |  |  |  |
| 1 | L lateral prefrontal & temporal | 72584 |  |  |  |  |  |
|  | L frontal operculum |  | 44 | -42 | 12 | 26 | .0066 |
|  | L IFG (pars orbitalis) |  | 47 | -42 | 30 | -6 | .0064 |
|  | L IFG (pars triangularis) |  | 45 | -50 | 20 | -2 | .0063 |
|  | L IFS/middle frontal gyrus |  | 44/45 | -46 | 22 | 24 | .0062 |
|  | L pMTG |  | 21 | -54 | -44 | -4 | .0039 |
|  | L precentral gyrus |  | 6 | -50 | -6 | 44 | .0033 |
|  | L mid superior temporal sulcus |  | 22 | -60 | -16 | -4 | .0032 |
| 2 | Medial frontal & cingulate | 23400 |  |  |  |  |  |
|  | dACC /SFG |  | 32 | -4 | 22 | 40 | .0062 |
| 3 | R IFG | 12032 |  |  |  |  |  |
|  | R IFG (pars orbitalis) |  | 47 | 38 | 21 | -10 | .0044 |

IFG = inferior frontal gyrus; IFS = inferior frontal sulcus; SFG = superior frontal gyrus; pMTG = posterior middle temporal gyrus; dACC = dorsal anterior cingulate cortex.

**Table 4:** ALE clusters for conjunction of young and older adults

| Cluster | Anatomical region | Volume (mm<sup>3</sup>) | BA | x | y | z | ALE value |
| --- | --- | --- | --- | --- | --- | --- | --- |
| 1 | L lateral prefrontal | 44960 |  |  |  |  |  |
|  | L IFG (pars orbitalis) |  | 47 | -42 | 30 | -6 | .0064 |
|  | L IFG (pars triangularis) |  | 45 | -50 | 20 | -2 | .0063 |
|  | L IFS/middle frontal gyrus |  | 44/45 | -46 | 22 | 24 | .0062 |
|  | L precentral gyrus |  | 6 | -42 | 6 | 40 | .0027 |
|  | L precentral gyrus |  | 6 | -46 | 0 | 44 | .0026 |
| 2 | L medial frontal & cingulate | 15664 |  |  |  |  |  |
|  | SFG/ dACC |  | 32 | -4 | 22 | 44 | .0060 |
|  | dACC |  | 24 | -4 | 26 | 36 | .0054 |
| 3 | R IFG | 5920 |  |  |  |  |  |
|  | R IFG (pars orbitalis) |  | 47 | 38 | 26 | -10 | .0044 |
| 4 | L pMTG | 2456 |  |  |  |  |  |
|  | L pMTG |  | 37 | -52 | -52 | -6 | .0030 |

IFG = inferior frontal gyrus; IFS = inferior frontal sulcus; SFG = superior frontal gyrus; pMTG = posterior middle temporal gyrus; dACC = dorsal anterior cingulate cortex.

This analysis performed three important functions. First it acted as a sanity check, indicating that the studies included in the meta-analysis did indeed identify activation in regions usually associated with semantic cognition (cf. Figure 1). Second, it highlighted areas that our analyses may *not* be sensitive to. We did not obtain significant ALE values in the vATL, which probably reflects well-known technical difficulties in acquiring signal from this region with fMRI, due to the proximity of air-filled sinuses (Devlin et al., 2000). This means that the studies in the meta-analysis are unlikely to be sensitive to potential age differences in the activation of this important semantic region. Finally, these analyses indicated that young and older individuals recruit broadly similar neural networks during semantic processing. This means that any age differences are relatively subtle in nature and should be interpreted in the context of a high degree of overall similarity.

### Contrasts of young and older adults

ALE maps derived from direct comparisons of young and older adults (all Y>O and O>Y peaks) are shown in Figure 3 (see Table 5 for details). Reduced activation in older participants was observed in a range of left-hemisphere regions linked with semantic processing, including a broad swathe of IFG extending into IFS, posterior temporal cortex including pMTG and ventral occipitotemporal regions, and a dIPC region extending into the intraparietal sulcus. Older adults also showed less activity in left hippocampus and in the occipital pole bilaterally. In contrast, enhanced activation for older individuals was most prominent in the right hemisphere, including the right IFG and a large area of right superior frontal and parietal cortex. Much of this right-hemisphere cluster overlapped with areas of the MDN (cf. Figure 1). The uncorrected maps also revealed smaller clusters of O>Y activity in left anterior IFS and in various regions of the DMN: pCing, vmPFC and bilateral vIPC (see Supplementary Figure 2).

**Figure 3:**
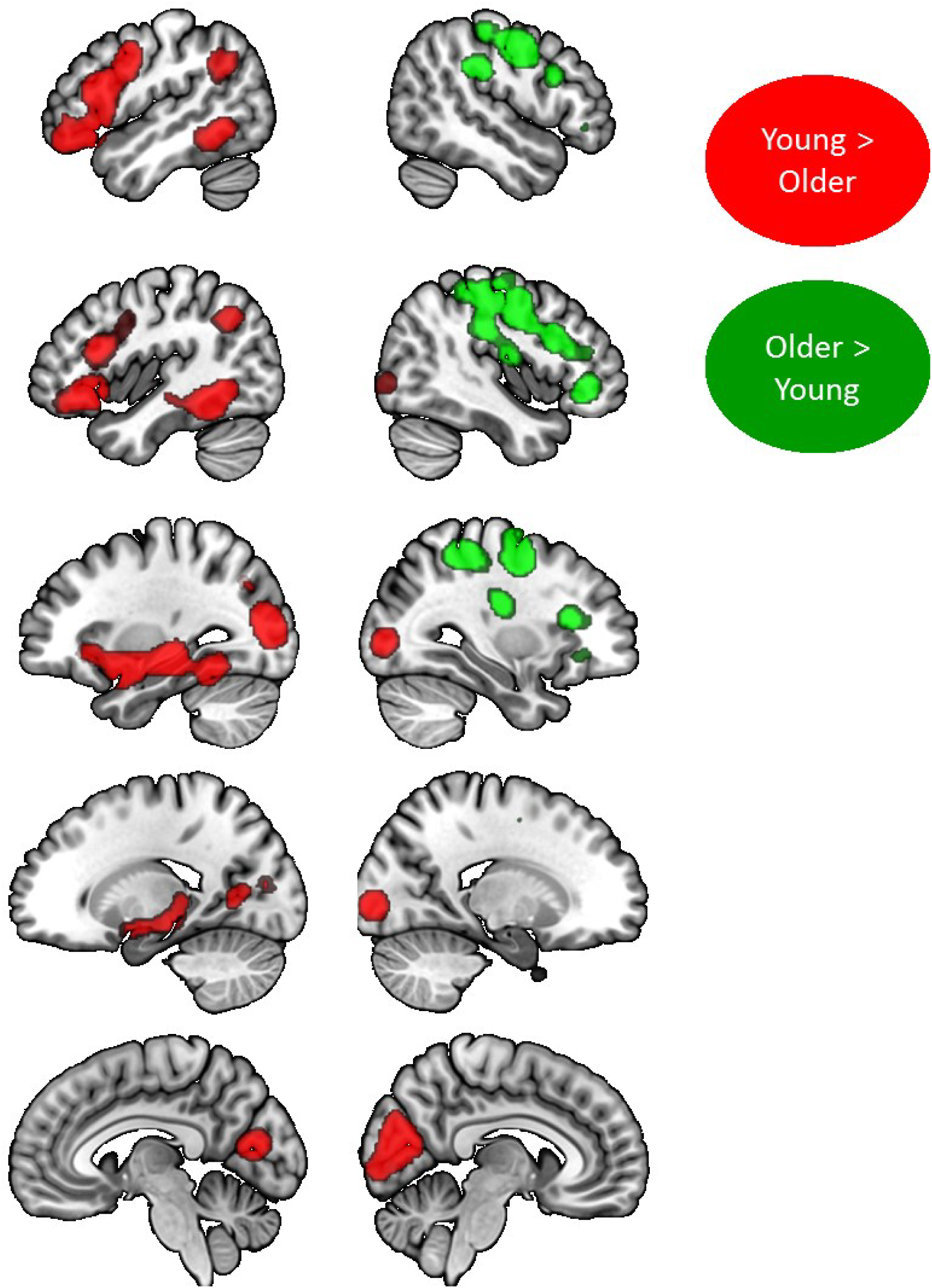
Activation likelihood maps for contrasts of young and older people. Results are presented at a threshold of *p* < 0.05, corrected for multiple comparisons at the cluster level.

**Table 5:**
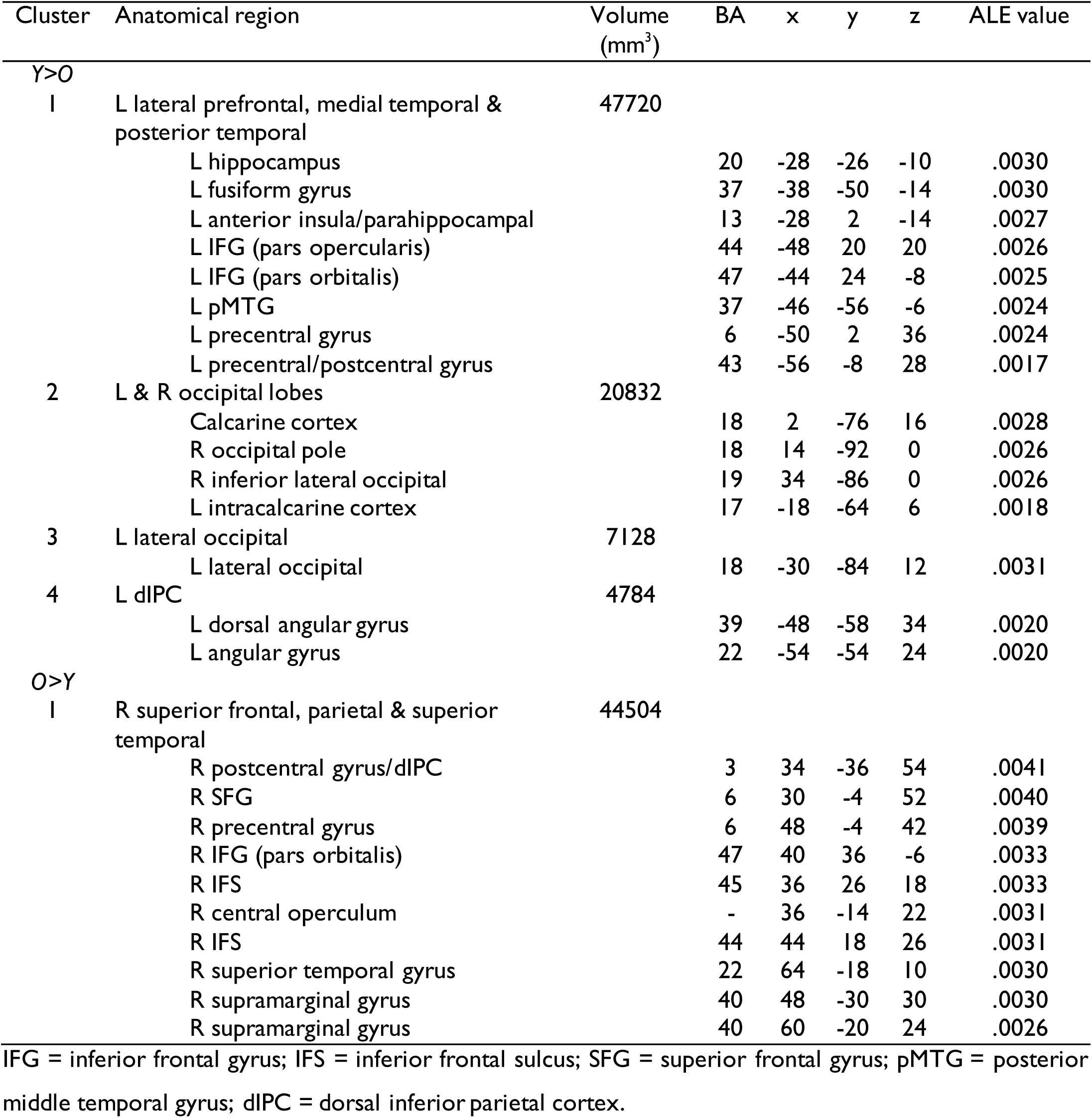
ALE clusters for Y>O and O>Y activation across all studies

Laterality analyses were performed to identify regions in which ALE values in one hemisphere were significantly higher than in the homologous region in the opposite hemisphere (see Figure 4). For the Y>O peaks, ALE values were significantly higher in the left hemisphere in areas of the precentral gyrus and parietal cortex, which overlapped with those identified in the main Y>O analysis. No regions exhibited higher activation likelihood in the right hemisphere, indicating that there was a clear leftward bias in Y>O peaks. The opposite was true for O>Y activations, with significantly higher ALE values in right IFG and right superior frontal and parietal cortex, relative to the analogous regions in the left hemisphere. The regions were also identified in the main O>Y analysis. Although this was an exploratory analysis (using an uncorrected threshold), its formal test of group by region interactions supports the above observation that older adults are more likely to show reduced activation in left-hemisphere regions and increased activation in the right hemisphere relative to the young.

**Figure 4:**
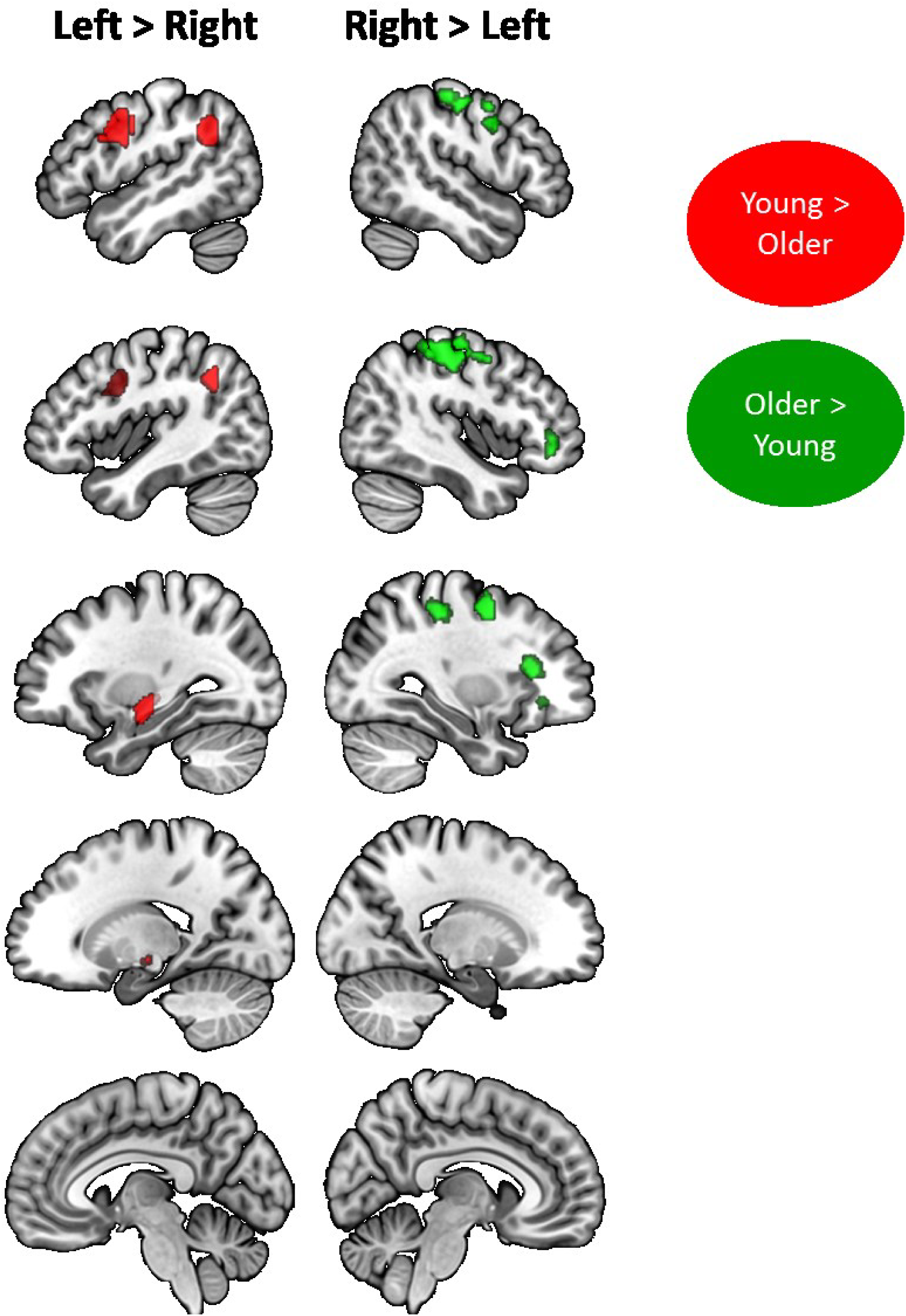
Laterality analysis of contrasts of young and older adults. Figure shows regions where ALE values were significantly higher in the left hemisphere compared with the homologous region in the right, and vice versa.

### Division of studies by behavioural effects

Here, we investigated whether the observed differences between age groups are related to behavioural performance differences between young and older participants, as predicted by theories of neurocognitive aging. We were able to compute an effect size for the performance difference between young and older adults in 29 of the 47 studies. The remaining studies either failed to report the relevant performance data or used covert or passive tasks with no behavioural measures. Older participants performed better than the young in 6 studies, while young participants outperformed the older participants in the rest (though in many cases the effect size was small and not statistically significant). We arranged the studies in order of effect size and used a median split to form them into two groups. The first, Performance-Equivalent group included the 6 studies with effect sizes favouring older individuals and other studies with smaller effects in favour of the young. The mean effect size in this set of studies was 0.06 (Cohen’s d), indicating that on average there was a negligible behavioural difference between the two age groups in these studies. The second, Performance-Reduced group included studies with larger effects favouring young people. The mean effect size in this set of studies was 1.01. This is a large effect in Cohen’s terminology and indicates that young people, on average, performed one standard deviation better than older people. The mean effect size differed significantly between the two sets of studies (*t*(30) = 4.74, *p* < 0.001). The mean ages of participants in the two sets of studies were very similar (young: 28.5 vs. 27.0 years; *t*(30) = 0.7, *p* = 0.50; older: 69.1 vs. 70.2 years; t(30) = 0.6, *p* = 0.55).

ALE maps for Y>O and O>Y peaks for these two subsets of studies are shown in Figure 5 (see Table 6 for co-ordinates). We regard these analyses as exploratory because there were a relatively small number of studies in each set. Therefore, we focus mainly on two sets of regions: those showing significant age effects in both Performance-Equivalent and Performance-Reduced studies, and those showing age effects in only one subset which overlapped with the results of the overall analysis. We first consider areas of reduced activation in older people. An important area of convergence across studies was in left IFG, which was under-activated by older people in both sets of studies, specifically in the ventral and anterior portions. Left medial temporal lobe/hippocampus also showed significantly reduced activation irrespective of performance differences. In other areas, effects appeared to depend on whether there were performance differences between the two groups. In some semantic regions, namely left pMTG and dIPC, reduced activation in older people only emerged in the Performance-Reduced studies. Thus, it appears that older people routinely activate left IFG to a lesser degree than young people, but that diminished activation in other key parts of the semantic network may only be seen when older people are performing at a lower level. In occipital cortices, the largest overlap of Y>O peaks with the main analysis was found for Performance-Equivalent studies.

**Figure 5:**
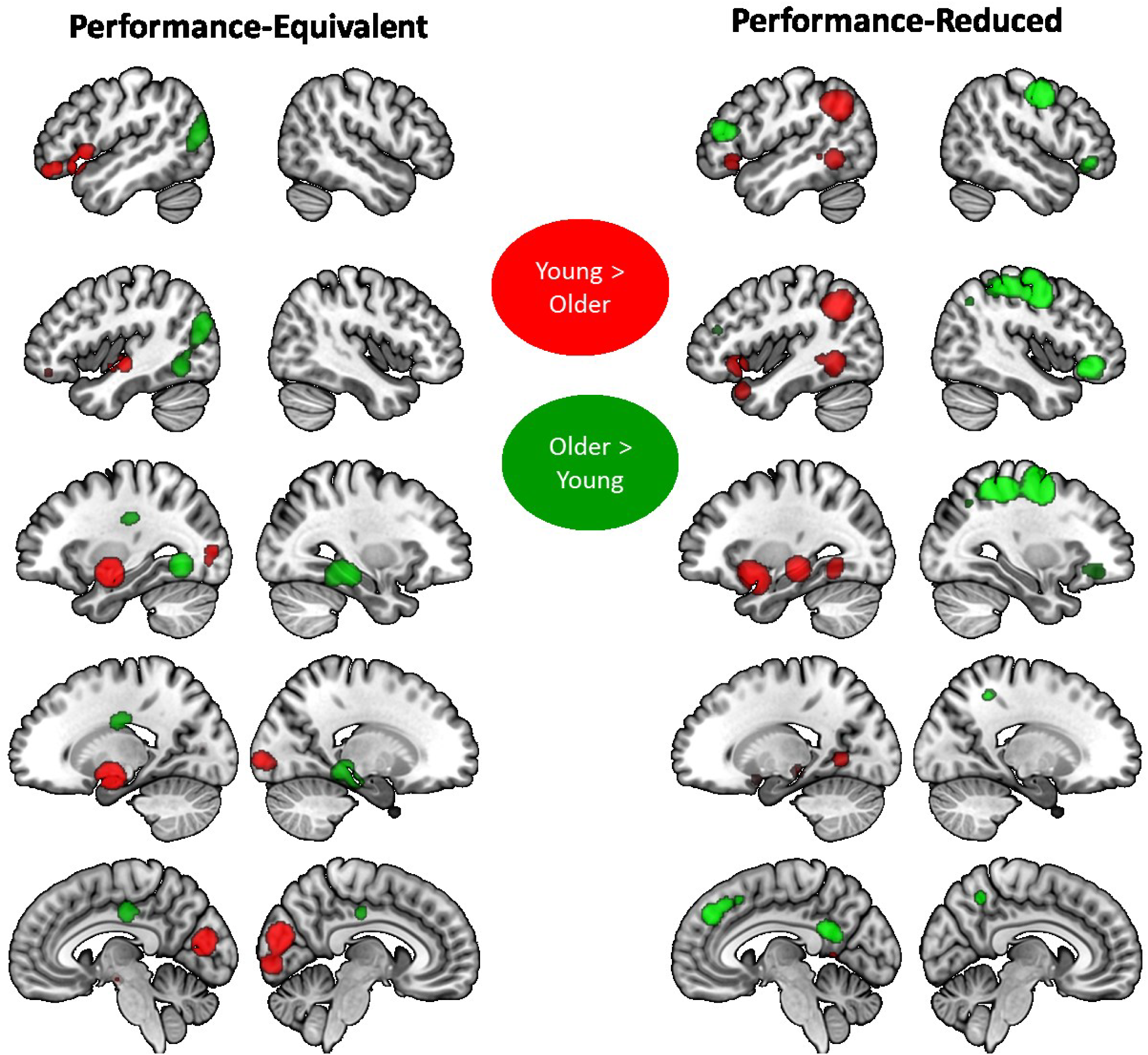
Activation likelihood maps for contrasts of young and older people, split by behavioural performance effects. Results are presented at a threshold of *p* < 0.05, corrected for multiple comparisons at the cluster level.

**Table 6:**
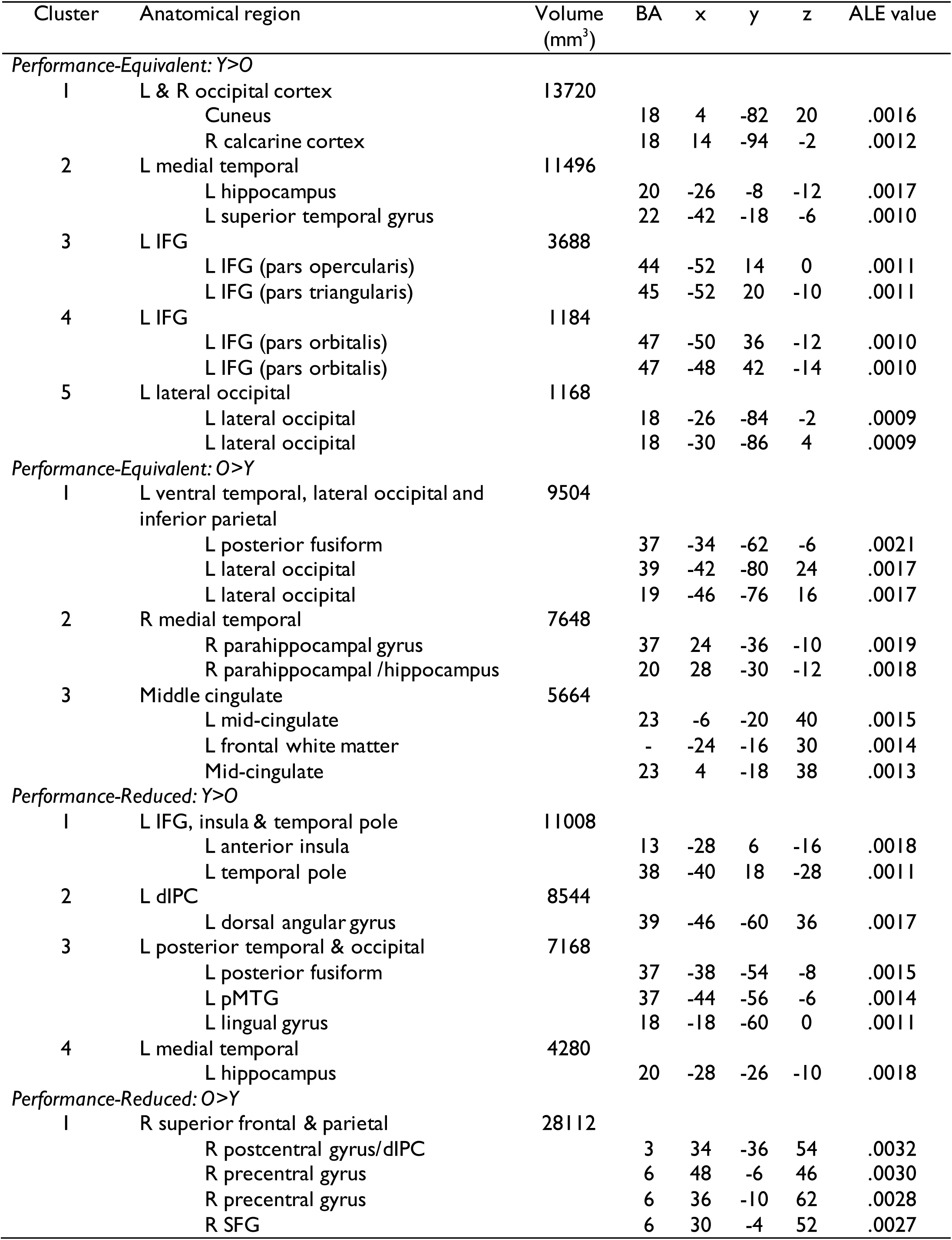
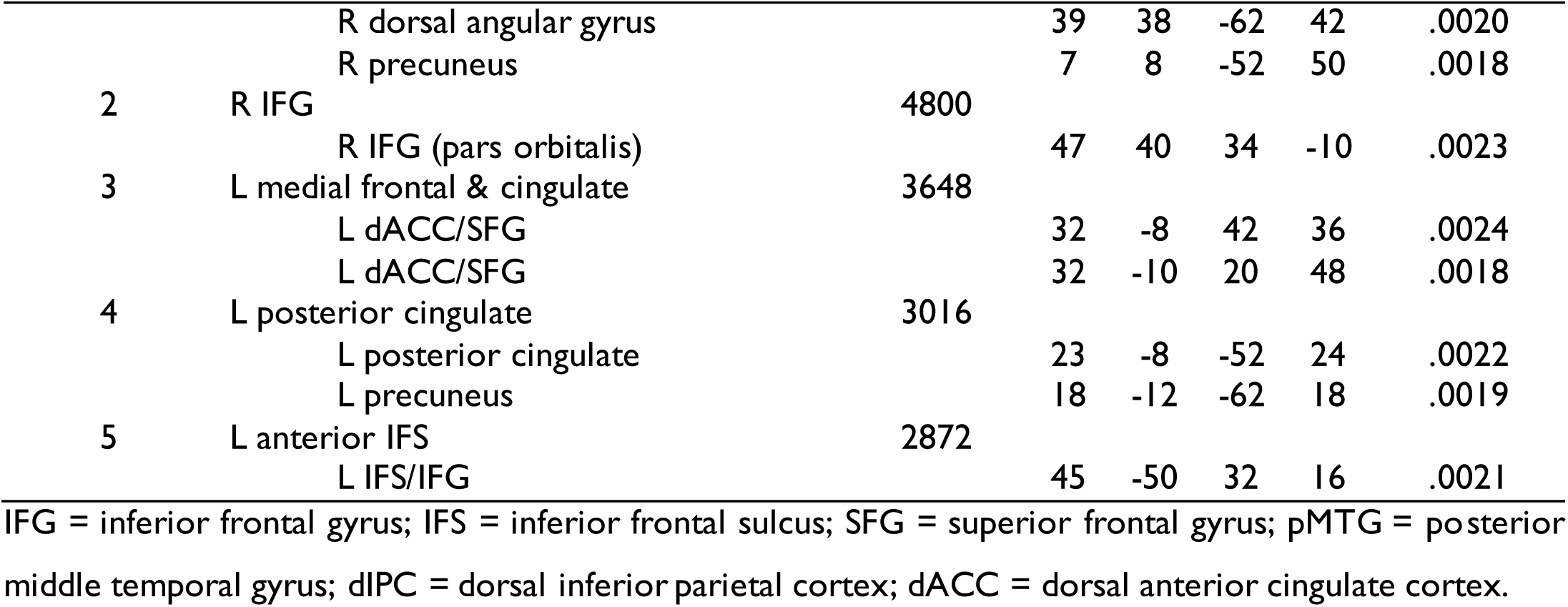
ALE clusters for Y>O and O>Y activation in Performance-Equivalent and Performance-Reduced studies

The O>Y contrasts showed minimal overlap across study type. For Performance-Reduced studies, there were large areas of activation in right frontal and parietal cortex, including right IFG, which corresponded closely to areas identified in the overall O>Y analysis. Other significant clusters were found in MDN regions: left anterior IFS (superior to the IFG region identified in Y>O analyses) and dACC (not found in the overall analysis). Thus, it appears that the tendency for older adults to increase activation in the right hemisphere and in the MDN tends to occur when they perform more poorly than young people. For Performance-Equivalent studies, older adults displayed significantly more activation in left vIPC and lateral occipital areas and in right medial temporal cortex. Two regions showed opposite direction age-related differences in the two subsets of studies: a region of left ventral occipitotemporal cortex showed reduced activation in older adults in Performance-Equivalent studies, and increased activation in the Performance-Reduced studies.

## Discussion

We used ALE meta-analysis of 47 functional neuroimaging studies to investigate age-related changes in the neural networks supporting semantic cognition. Separate analyses of young and older participants revealed that both age groups activated similar, left-lateralised networks, which included lateral prefrontal, medial frontal and posterior temporal regions. Against this backdrop of broad similarity, however, there were a number of areas in which recruitment varied as a function of age. Older people demonstrated less activation in left-hemisphere regions associated with control and regulation of semantic processing (IFG, pMTG, dIPC). In pMTG and dIPC, age-related differences were only robust for studies in which older people performed more poorly than their younger counterparts, while in left IFG, older and younger adults differed regardless of relative performance levels. Older people also showed decreased activation of occipital cortex, which appeared to be driven mainly by studies in which the groups performed at an equivalent level. In other areas, older people demonstrated more activation than the young. These encompassed right frontal and superior parietal lobes, including right IFG and areas of the MDN. This increased activation appeared to be driven mainly by studies where older people performed at a lower level than young people. Taken together, these findings indicate a shift from the left-lateralised semantic network in later life, with less activation in left-hemisphere regions linked specifically with semantic processing and greater activity in the right hemisphere and in elements of the MDN. The most prominent changes seemed to occur when older adults were unable to maintain task performance at the same level as young people. Here, we consider the extent to which these findings are compatible with existing theories of neurocognitive ageing and where they provide new evidence for age-related differences that may be specific to semantic cognition.

In the Introduction we outlined two leading theories of neurocognitive ageing which propose that there are large-scale age-related shifts in patterns of brain activity. The PASA theory (Davis et al., 2008; Dennis and Cabeza, 2008; Grady et al., 1994) holds that older adults are less efficient at processing visual stimuli and therefore exhibit reduced activation in posterior occipital and temporal regions. To compensate for this decline, older individuals are proposed to upregulate activation in prefrontal regions associated with executive control. We found support for the first prediction of this theory: across all studies, there were age-related activation reductions in primary visual cortex and in left ventral occipitotemporal regions associated with visual word and object recognition. However, the data were not unambiguous, as there were also smaller age-related activation *increases* in other occipital and fusiform regions. The evidence for increases in recruitment of prefrontal regions was more mixed. Contrary to PASA, left IFG – a major site for regulation of semantic processing – was reliably *less* active in older people, though they did show more activation in a large swathe of right PFC. This suggests that, where semantic processing is concerned, different areas of PFC are affected by ageing in different ways, consistent with findings from studies of episodic and working memory (Rajah and D’esposito, 2005). Our results are inconsistent with a general picture of a posterior-to-anterior shift, at least in the studies of semantic cognition surveyed.

HAROLD takes the view that cognitive ageing is associated with reductions in the asymmetry of activation patterns, particularly in the prefrontal cortices (Cabeza, 2002; Grady, 2012). Consistent with previous meta-analyses (Binder et al., 2009; Noonan et al., 2013), we found that young participants recruited a strongly left-lateralised network during semantic tasks. A reduction in lateralisation in the semantic domain would therefore entail less left-hemisphere and more right-hemisphere activity in later life. This is what we observed, broadly speaking. Older adults reliably demonstrated less left-hemisphere and more right-hemisphere activation. These large clusters included left and right IFG. The direct analysis of age-related differences in lateralisation confirmed the presence of localised effects in IFG, and also implicated other frontal and parietal sites.

One view of HAROLD is that more bilateral recruitment of neural resources is a compensatory effect that helps to maintain performance in older age (Cabeza, 2002). In young people, greater right IFG activation is observed for highly demanding semantic tasks that require more executive control (Krieger-Redwood et al., 2015; Noonan et al., 2013). Upregulation of the activation of this area in older people may therefore reflect increased reliance on this demand-related mechanism. We found that older adults’ additional activation in right IFG (and elsewhere in the right hemisphere) was most robust in studies where they performed more poorly than young. One interpretation is that these studies employed tasks that older participants found more difficult and which therefore elicited greater recruitment of right IFG to support performance to some degree (although not to the level of the young participants). Of course, another possibility is that right IFG upregulation does not support performance in older people, and may be a cause of rather than a response to performance declines. This debate is unlikely to be fully resolved by the correlational methodology of neuroimaging studies alone, particularly by cross-sectional studies, which also struggle to establish interpretable associations between age effects and performance (see Morcom and Johnson, 2015). TMS studies, however, permit assessment of the effects of temporary disruption of function, and one such study comparing young and older adults provided some support for the view that dorsolateral prefrontal regions contributing to performance are more bilateral in older age, at least during episodic memory retrieval (Rossi et al., 2004). No TMS studies to date have investigated such effects on semantic cognition specifically. It worth noting that increased right prefrontal recruitment is frequently observed in aphasic patients following stroke and is associated with better recovery of language function, at least in some cases (Saur et al., 2006; Winhuisen et al., 2005).

One possible prediction of a compensatory account was an upregulation of left IFG in older people, as well as in right IFG. Young people show reliable increases in left IFG activation for more demanding semantic tasks (Badre et al., 2005; Krieger-Redwood et al., 2015; Noonan et al., 2013). However, we found that older individuals activated left IFG less than the young. A potential explanation for this discrepancy is offered by the CRUNCH theory outlined in the Introduction (Reuter-Lorenz and Cappell, 2008). On this view, older people can successfully maintain their performance up to a point by increasing activation of task-related brain areas above that of their younger peers. Under more demanding conditions, however, this effect reaches a plateau beyond which activation in older people tails off, and young people show greater activation. Since left IFG is a core element of the semantic network in young people, and demonstrates robust activation across all semantic tasks, it may have little spare capacity for additional recruitment in older age. In order to address this hypothesis in detail, direct manipulation of task demand in people of different ages will be needed (Reuter-Lorenz and Cappell, 2008; Schneider-Garces et al., 2010).

Finally, many of the results of the present meta-analysis are consistent with the idea that activation shifts in later life away from neurally specialised regions and towards more task-general areas. In the current study, older adults displayed reduced activity in several core areas of the left-hemisphere semantic network, including left IFG, pMTG and dIPC. These areas have been linked particularly with executive regulation of semantic knowledge (Jefferies, 2013). This finding may indicate reduced efficiency of such processes in older age, in line with executive control declines in working memory and episodic memory tasks (McCabe et al., 2010). Although behavioural studies frequently find little effect of healthy ageing on semantic tasks (Nilsson, 2003; Nyberg et al., 1996; Park et al., 2002; Rönnlund et al., 2005; Salthouse, 2004; Verhaeghen, 2003), in the current meta-analysis, older people often performed more poorly than the young. One reason for this discrepancy may be that tasks used in neuroimaging studies rely more heavily on controlled processing (e.g., fluency or semantic association judgements), in contrast to behavioural studies that typically employ measures of vocabulary size. Studies of ageing rarely use tasks that manipulate semantic control specifically (e.g., Badre et al., 2005). Such studies will help to determine whether semantic impairments in older age, where these do occur, are related to poorer control.

At the same time, we observed reliable age-related increases in activation in areas of the domain-general MDN, including right IFS and middle frontal gyrus, right dIPC and dACC. This may indicate that older people draw more heavily on flexible domain-general processing resources to compensate for under-activation of the core semantic network, i.e., the additional recruitment of MDN regions reflects neurocognitive flexibility, as articulated by Lövdén et al., (2010). As noted in the Introduction, our data cannot determine whether this additional recruitment actually benefits performance, or is secondary to a reduction in the specificity of neural responses (Grady, 2012; Grady et al., 1994; Li et al., 2001; Logan et al., 2002). However, like the CRUNCH hypothesis, the neurocognitive flexibility theory of additional recruitment makes other testable predictions. Additional engagement of the MDN in older adults should be found across task domains (e.g., semantic cognition and episodic memory), and should depend on task demand, so that engagement of domain-general regions and networks in older people at low demand should resemble those in young people at high demand.

We found no strong evidence for age-related differences in DMN activity (although the direction of findings at the more lenient uncorrected threshold was for several small areas of reduced activity in older people). The role of DMN regions in semantic cognition is currently unclear, with some researchers arguing that some regions classified as within the DMN (vIPC and pCing, in particular) make important contributions to semantic processing (Binder and Desai, 2011). Others claim that there is a strong distinction between the DMN and the semantic network (Humphreys et al., 2015). However, evidence from one previous fMRI study suggests that DMN activation is negatively associated with semantic performance. Persson et al. (2007) compared young and older participants performing verb generation, a difficult semantic task. They found that DMN regions (particularly pCing) were deactivated by the semantic task and that the degree of deactivation increased with increasing task demand. Older adults exhibited less deactivation than young people in the most demanding conditions and, importantly, individuals with greater deactivation in right posterior cingulate displayed better task performance. This study suggests that increased DMN activation in older people reflects a failure of older adults to deactivate this network, which may in turn have negative effects on semantic performance. This is an important possibility for future studies to consider, in light of increasing evidence for interaction between DMN and semantic regions (Vatansever et al., 2017).

### Convergence with previous meta-analyses

The results of the present meta-analysis are broadly consistent with previous meta-analyses that have investigated more general effects of healthy ageing on functional brain activity (Li et al., 2015; Spreng et al., 2010). These meta-analyses included many of the studies we investigated but also included numerous studies of episodic and working memory, perception and executive function that fell outside our more targeted approach. Similar findings of age-related reductions in activation of visual cortices were reported by both Spreng et al. (2010) and Li et al. (2015) and may be a consequence of impaired or less differentiated visual processing in later life. Our data also revealed greater activation with age in other visual regions, which may be consistent with a dedifferentiation view, i.e. that visual cortical function is less specific rather than simply impaired (Carp et al., 2011; Park et al., 2004). Reduced activity in the left hippocampus was also reported in both previous meta-analyses as well as the present study, and may reflect reductions in the frequency with which this region is engaged in incidental encoding of novel experiences into episodic memory (Daselaar et al., 2003). Likewise, both previous meta-analyses found that older adults demonstrated reduced activity in areas of left IFG, consistent with our findings. In contrast, a meta-analysis of subsequent memory effects in episodic memory studies found no differences between young and older people in left IFG or hippocampus (Maillet and Rajah, 2014). This result does not conflict with our findings; older adults may be less likely to engage this region in semantic processing but when they do engage it, the activation is just as effective for episodic memory encoding as in the young (Maillet and Rajah, 2014; Morcom et al., 2003).

A difference between our results and those of the two earlier more inclusive meta-analyses (Li et al., 2015; Spreng et al., 2010) is that they found additional recruitment by older people of more posterior left PFC regions, which we did not. In addition, neither previous study found evidence for reduced activation of left pMTG or dIPC. It is likely that our analysis had greater power to detect such effects as a consequence of focusing specifically on semantic tasks that provide strong activation in these regions. Increases in right PFC regions were also found in previous meta-analyses, particularly when older people performed more poorly than young. More generally, both previous meta-analyses reported increased activation in older participants in MDN regions, which accords with our findings. In summary, many of the age-related differences we found were consistent with those reported for other cognitive domains, though we also found some additional age-related differences. Direct comparisons in future studies will be able to establish whether these differences are specific to semantic cognition.

### Implications for future studies

Meta-analyses can be useful not only in synthesising the current state of knowledge in a domain but also in plotting where the limits of our current understanding lie. This meta-analysis has identified two lacunae in our understanding of age-related changes in semantic cognition. First, we note that the literature is heavily biased towards verbal semantic processing. Forty of the analysed studies either used lexical stimuli or required verbal responses, while only 13 presented non-verbal stimuli (usually pictures but in two studies, smells). This is important because non-verbal semantic processing, in addition to being an essential part of everyday life, engages a different distribution of brain regions to verbal semantic cognition. While verbal semantic processing is strongly left-lateralised (particularly for written words), non-verbal stimuli elicit more bilateral patterns of activation (Krieger-Redwood et al., 2015; Rice et al., 2015a; Rice et al., 2015b; Visser et al., 2010). As a consequence, the general shift in activation away from left-hemisphere regions and towards contralateral activation may be less prominent for non-verbal processing. The degree to which the present findings apply to non-verbal processing therefore remains an open question, as does the status of non-verbal semantic cognition in ageing more generally. However, one simple prediction of the neurocognitive flexibility theory of additional recruitment, consistent with our data for predominantly verbal studies, is that older people will show greater activation of MDN regions than young people in non-verbal semantic tasks.

Second, the studies included in this meta-analysis did not consistently report activation even in young people in the vATLs, which are now known to be a key region in the representation of semantic knowledge (Binder and Desai, 2011; Humphreys et al., 2015; Patterson et al., 2007). The failure to detect engagement of this area most likely reflects a combination of methodological factors that reduce the likelihood of activity in this area being sampled properly (Visser et al., 2010). These include poor signal in the vATL in fMRI studies, due to the proximity of air-filled sinuses (Devlin et al., 2000), its extreme ventral position in the brain which can lead to it being excluded from image acquisition (Visser et al., 2010) and the use of resting baselines that do not adequately control for task-unrelated semantic processing (Humphreys et al., 2015). When these issues are addressed, semantic cognition does reliably activate this area (e.g., Hoffman et al., 2015; Humphreys et al., 2015; Spitsyna et al., 2006). However, since vATL was not reliably activated in the studies included in the meta-analysis, we are unable to draw any conclusions about possible age effects in this region. This is an important target for future work, because vATL and left IFG are thought to play complementary roles in semantic task performance (Lambon Ralph et al., 2017).

## Acknowledgements

PH was funded by The University of Edinburgh Centre for Cognitive Ageing and Cognitive Epidemiology, part of the cross council Lifelong Health and Wellbeing Initiative (MR/K026992/1). Funding from the Biotechnology and Biological Sciences Research Council (BBSRC) and Medical Research Council (MRC) is gratefully acknowledged. AM is a member of CCACE.

---

**Supplementary Figure 1:**
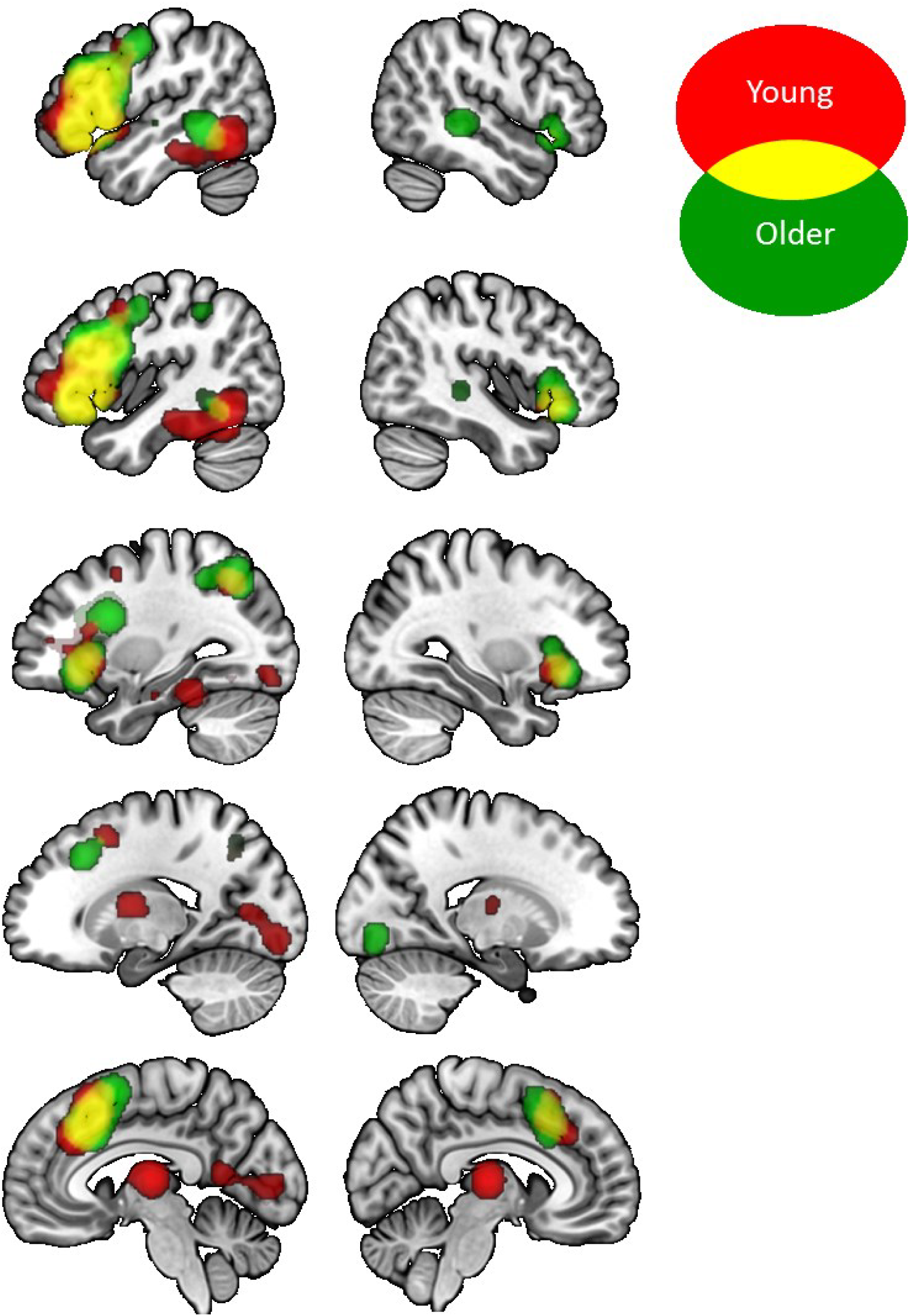
Activation likelihood maps for separate analyses of young and older people, at an uncorrected statistical threshold. Results are presented at an uncorrected voxel-height threshold of *p* < 0.01 with an extent threshold of 1000mm^3^.

**Supplementary Figure 2:**
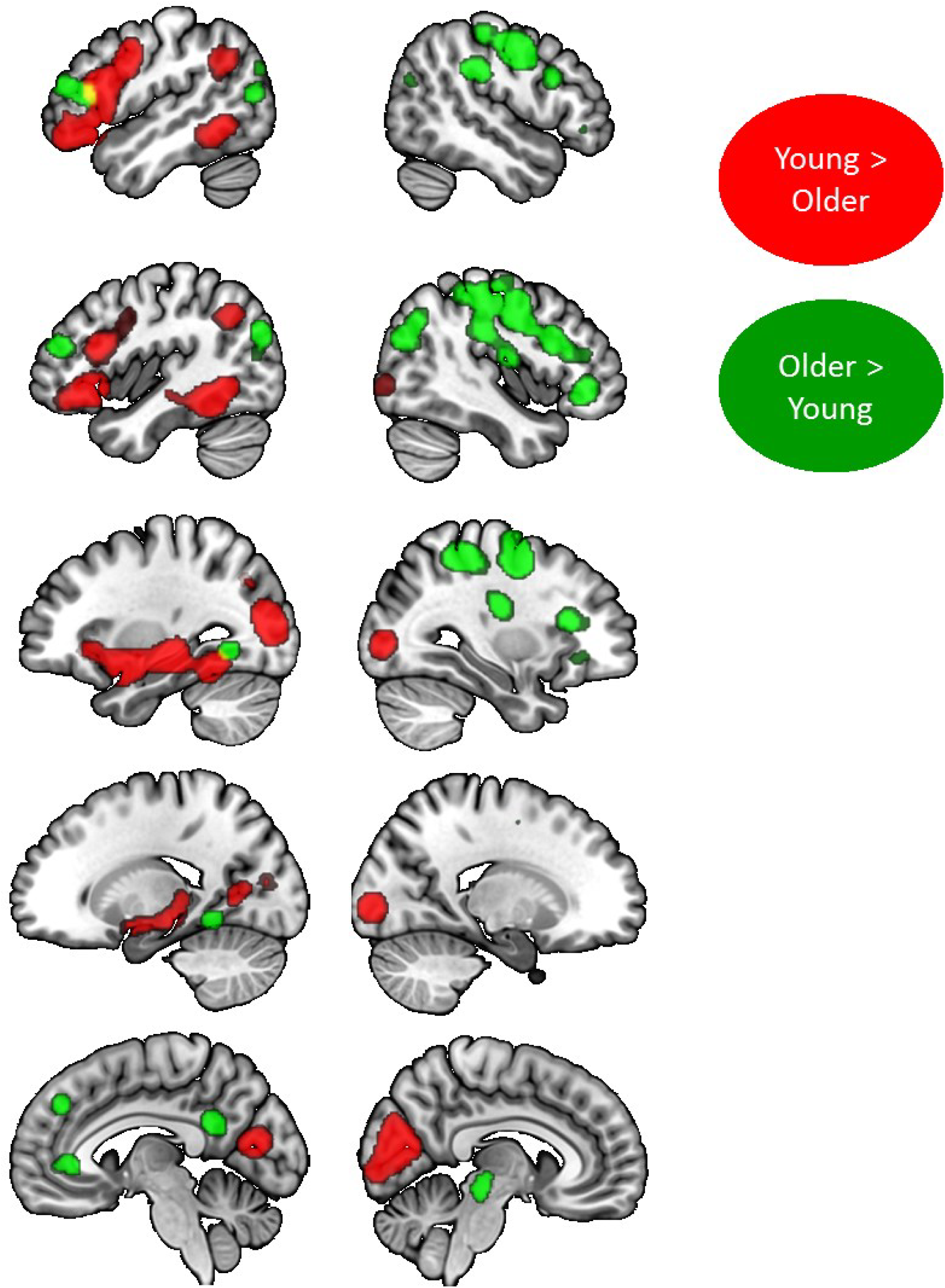
Activation likelihood maps for contrasts of young and older people, at an uncorrected statistical threshold. Results are presented at an uncorrected voxel-height threshold of *p* < 0.01 with an extent threshold of 1000mm^3^.

